# Patient-specific computational mechanics of functional lumbar spine units

**DOI:** 10.64898/2026.06.03.729850

**Authors:** Ivan Fumagalli, Michela Campioni, Anna Sirtori, Stefano Pagani, Riccardo Levi, Letterio Salvatore Politi, Gabriele Capo, Paola F. Antonietti

## Abstract

In the current clinical practice, the diagnosis of spinal disorders and their surgical planning are critically based on imaging data. To complement this data, patient-specific finite element models have been developed and showed to be powerful tools for evaluating spine mechanics. Most of them rely on Computational Tomography (CT) scans – which have a high resolution but are seldom available in routine clinical practice – while only a recent few models are on less invasive Magnetic Resonance Imaging (MRI). Yet, despite the proliferation of these computational models, encompassing detailed anatomical and functional information, the rheological assumptions they are built upon are based on tissue-sample mechanical response data, which leaves a gap in the quantitative analysis on how such assumptions influence the macroscopic response of a functional spinal unit. Aiming at addressing these shortcomings, the main purpose of this work is to introduce a quantitative computational assessment of the macroscopic impact of commonly adopted rheological models – from linear elasticity to fiber-reinforced nonlinear hyperelasticity – in several loading conditions, focusing on a lumbar unit which is considered as a typical benchmark system. We also propose a reconstruction procedure to accurately describe subject-specific anatomy from MRI data, including the intervertebral disc and its nucleus pulposus. Bones are modeled as linear elastic media, whereas for the AF, we consider three different mechanical models – namely, isotropic linear elasticity and the Holzapfel-Gasser-Ogden model with and without fiber reinforcement. Model verification on an idealized geometry demonstrates numerical consistency, while parametric orthostatic simulations highlight the need for nonlinear formulations to capture anisotropy and strain-stiffening behavior of the intervertebral disc. Then, we carry out flexion, lateral bending, and torsion tests on a subject-specific reconstructed functional unit, for which we provide parametric analysis in terms of momentum magnitude and resulting range of motion. These tests further confirm the need for a nonlinear rheology of the annulus fibrosus and provide a quantitative assessment of the differences between the constitutive laws considered. Moreover, successful comparisons with the literature, in terms of macroscopic deformation under several loading conditions, serve as partial validation for our computational model.

## 1 Introduction

Low back pain is one of the leading causes of disability worldwide and, together with other spinal disorders, has a huge impact on patients and society as a whole, due to the associated lifelong care and loss of workforce [1, 2]. For severe cases of degenerative disc disease, spondylosis, and disc herniation – which are the most common and debilitating spine disorders – the only clinical treatment is surgical intervention, such as spinal fusion or microdiscectomy, since physical therapy and anti-inflammatory drugs can only act as conservative measures [3]. The success of surgical treatment is highly dependent on diagnosis, surgical planning, and rehabilitation, which are all based on diagnostic imaging data (typically MRI and/or X-ray scans). In this respect, computational modeling has recently evolved as an increasingly effective tool to complement clinical practice with patient-specific in-silico models, capable of capturing additional details of the patient’s state and anticipating treatment response in different intraoperative and post-intervention scenarios [4, 5, 6].

Due to the anatomical complexity of the spine and the varied loading conditions it sustains, both under physiological and pathological conditions, several models of the spine have been proposed in the computational literature. Most of them are based on Finite Element (FE) analysis, yet they differ in the level of detail in anatomical features, mechanical rheology, and loading-condition complexity. Models encompassing the entire thoraco-lumbar section or the totality of the spine – possibly including the rib cage and other torso elements – typically rely on a parametrization of the overall structures and simplified geometry and/or mechanical response for individual bones and ligaments [7, 8, 9, 10, 11]. El Borajarami et al.[12], on the other hand, construct and validate a model of the full spine and the main spinal erectors based on MRI scans of a healthy subject, thus improving the anatomical detail at the cost of reduced flexibility to account for pathological conditions or inter-patient variability. Due to the computational effort that it entails, image-based geometry reconstruction is used more extensively in computational models focused on a portion of the spine. In this respect, the lumbar tract is typically considered the benchmark framework for validating computational models. For example, eight lumbar models are assessed in physiological conditions by Dreischarf et al.[13], compared to macroscopic deformation measurements obtained in vivo, while other works study the sensitivity of the FE model with respect to mechanical parameters under different loadings [14, 15, 16].

In-silico models of the spine proposed in the literature also differ in the rheological assumptions they make, especially about soft tissues. Individual bones are typically modeled as rigid or linear elastic bodies [13, 12], possibly with orthotropic response in some regions to account for trabeculation [17, 9], while cartilaginous ligaments are always considered deformable, with either a linear or Ogden-type response [9]. The innermost nucleus pulposus of the intervertebral disc (IVD) is uniformly treated as an incompressible fluid-filled cavity, with very few exceptions adopting linear elasticity [18, 19], whereas for the annulus fibrosus (AF), several mechanical models are considered in the literature. They range from linear elasticity [12, 20] to nonlinear poroelasticity [21, 22] and to hyperelastic rheologies such as the Mooney-Rivlin [23, 24, 13, 25, 26] and the Holzapfel-Gasser-Ogden models [27, 28, 9, 14]. Almost all AF models also account for collagen fibers, either by anisotropic terms in the tissue’s mechanical energy or by additional layers of fibrous materials. However, to the best of the authors’ knowledge, no quantitative analysis has been reported on the impact of fiber reinforcement on macroscopic spine deformations under different loads.

Regarding geometry reconstruction, most of the aforementioned image-based studies are based on CT scans, due to their typically high spatial resolution. For these data, advanced and automated analysis techniques and mesh generation pipelines are available and continuously improved, often exploiting the efficiency of artificial-intelligence tools [29, 30, 31, 22, 32, 33]. However, CT scans are rarely available in routine clinical practice, where less invasive MRI acquisitions are typically collected. Indeed, MRI is better suited to assess functional information on the patient’s condition and is thus the reference imaging data for diagnosing pathological conditions and planning intervention when surgery is required. For this reason, some recent works focused on anatomical reconstruction from MRI scans, addressing the main challenges of this type of data [34, 35]. For a broader review of both reconstruction techniques and computational modeling of the thoraco-lumbar spine, we refer to Carpenedo et al.[36].

In the framework described above, this work aims to understand and assess some of the most widely adopted modeling choices – typically motivated solely by anatomical considerations – in terms of their macroscopic effects on a lumbar unit. For this purpose, we consider real patient-specific vertebral geometries segmented from volumetric MRI acquisitions, with a physiological disc reconstructed from established literature data. In this computational domain, we consider linear elastic bones, while for the AF we compare three different mechanical models – namely isotropic linear elasticity, isotropic nonlinear neo-Hookean hyperelasticity, and the Holzapfel-Gasser-Ogden model with fiber reinforcement – in several loading conditions, to quantify the effects of nonlinear mechanical response and anisotropy.

The structure of the work is as follows. In Section 2 we describe the adopted mathematical models, with particular focus on the AF, and introduce its numerical discretization and the imposition of loading conditions. Section 3 illustrates the procedure for MRI-based geometry reconstruction and mesh generation. Verification tests for the numerical model on a simplified geometry are reported in Section 4. Section 5 presents the main results for the reconstructed subject-specific geometry, and compares the outputs of the different mechanical models considered. Finally, Section 6 reports the conclusions of the work and provide directions for further development.

## 2 Mathematical models and numerical methods

We consider the Functional Spinal Unit (FSU) depicted in Fig. 1, composed of the two lumbar vertebrae L3 and L4 and the corresponding AF of the intervertebral disc. The disc’s nucleus pulposus (NP) is treated as a fluid-filled cavity in the FSU geometry. The whole FSU is taken as a deformable solid – with nonhomogeneous mechanical properties – whose current configuration ℬ is obtained from a reference configuration ℬ^*\**^ via a map ***χ*** : ℬ^*\**^ *→* ℝ^3^, namely each point **x** *∈ ℬ* is the image **x** = ***χ*** (**X**) of a material point **X** *∈ ℬ*^*\**^. We assume that the reference configuration is regular and that the map ***χ*** is invertible and piecewise smooth in each component of the FSU (i.e. in each vertebra, the AF, and each facet joint). Moreover, we denote by **u**: ℬ^*\**^*→* ℝ^3^ the deformation field, such as ***χ*** (**X**) = **X** + **u**(**X**). The boundary of the reference configuration is partitioned into a clamped portion 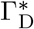, corresponding to the inferior plate of L4, and the rest 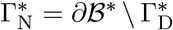, which includes the boundary of the nucleus pulposus, as illustrated in Fig. 1.

**Figure 1:**
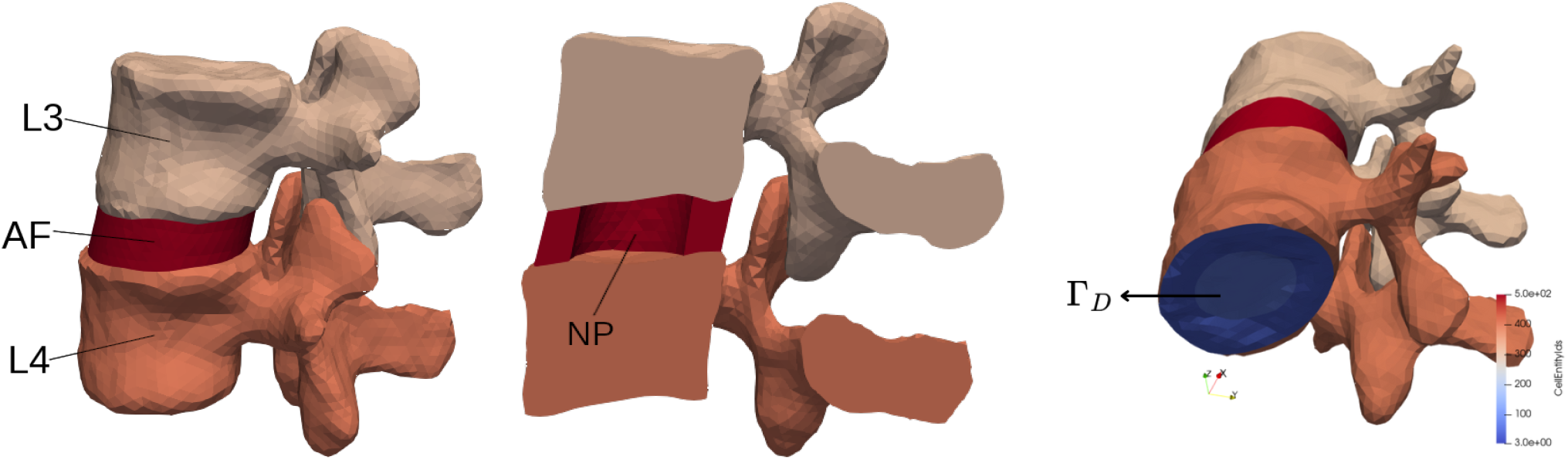
Domain ℬ^*\**^ with the functional spinal unit elements (left), the nucleus pulposus cavity (NP, center), and the Dirichlet boundary 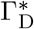 (right).

The deformation **u** is determined by the following equations of mechanical equilibrium:

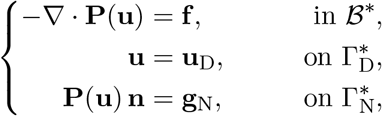

where **P**(**u**) is the first Piola-Kirchhoff stress tensor, **f** the external body force per unit volume (e.g., gravity or distributed load), **u**_D_ a prescribed deformation on 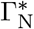, and **g**_N_ a stress distribution applied to 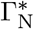.

The system is closed by the constitutive relations described below, with all model parameters summarized in Table 1. In particular, a uniform linear elastic rheology is adopted in the vertebral bodies, while different choices are considered for the AF, thus defining the scenarios described in Table 2.

**Table 1:**
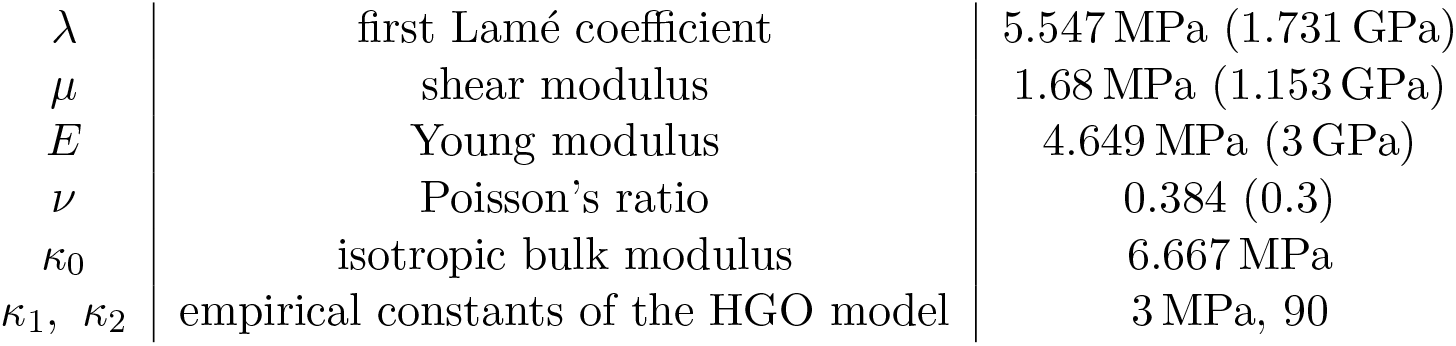
Physical parameters of the model. The values in the parentheses are employed in the vertebral bodies, while the others refer to the AF.

|  |  |  |
| --- | --- | --- |
| $\lambda$ | first Lamé coefficient | 5.547 MPa (1.731 GPa) |
| $\mu$ | shear modulus | 1.68 MPa (1.153 GPa) |
| $E$ | Young modulus | 4.649 MPa (3 GPa) |
| $\nu$ | Poisson's ratio | 0.384 (0.3) |
| $\kappa_0$ | isotropic bulk modulus | 6.667 MPa |
| $\kappa_1, \kappa_2$ | empirical constants of the HGO model | 3 MPa, 90 |

**Table 2:**
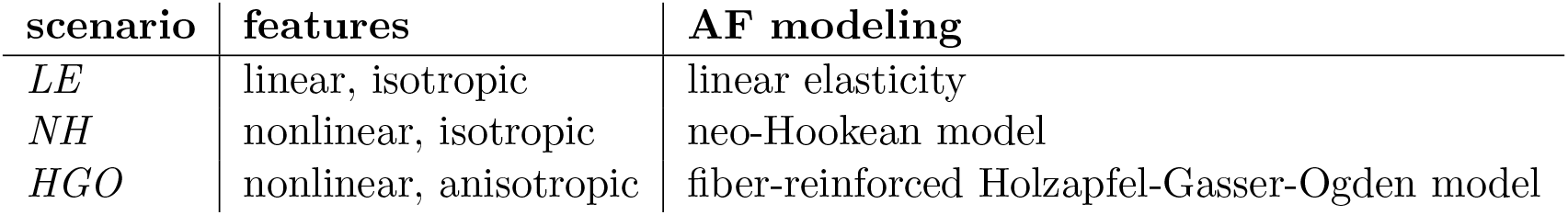
Definition of scenarios depending on the modeling of the AF. The vertebral bodies are always modeled as linear elastic.

### 2.1 Constitutive relations

The constitutive relation used for the vertebral bodies and for the AF in the *LE* scenario is that of linearly elastic, homogeneous, and isotropic materials, which reads as follows:

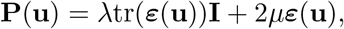

where **I** is the identity tensor and ***ε***(**u**) is the strain tensor 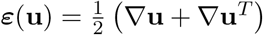. The Lamé coefficients *λ* and *µ* are typically obtained by conversion of the measurable Young modulus *E* and Poisson’s ratio *v*, as follows:

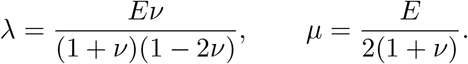

In the aforementioned scenarios *NH* and *HGO*, instead, we consider hyperelastic constitutive equations for the AF. We denote by **F** = **I** + ∇**u** = ∇***χ*** the deformation gradient, by **C** = **F**^*T*^ **F** the right Cauchy-Green tensor, and by *ψ* the hyperelastic energy, such that 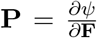. The deformation gradient can be split into a volumetric component **F**_vol_ = *J* ^1*/*3^**I** – where *J* = det (**F**) – and an isochoric part 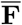, such that det 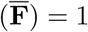 and 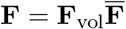. Correspondingly, we denote by 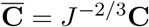 the isochoric part of **C**. Building upon these definitions, the neo-Hooke model employed in scenario *NH* is characterized by the following homogenous and isotropic energy:

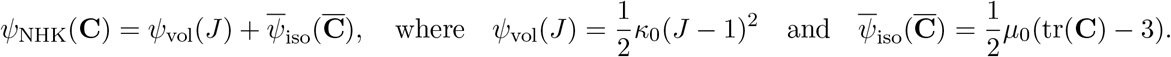

Here, *µ*_0_ = 2*C*_10_ denotes the initial shear modulus of the neo-Hookean model, and should not be confused with the Lamé coefficient *µ* introduced in equation (2.1) for the linear elastic scenario; the two parameters coincide only in the small strain limit.

The Holzapfel-Gasser-Ogden model [37, 28], on the other hand, also entails fiber orientation, described by two orthogonal unit vectors **a**_04_ and **a**_06_ corresponding to the local main fiber direction and the local fiber sheet normal direction, respectively. In accordance with the work by Bellina et al.[9], we define the sheet normal **a**_06_ as directed radially, while the fiber direction creates an angle of −33° with the AF axis at the boundary with the nucleus pulposus, and progressively rotates until it reaches an angle of +33° at the external boundary. Further details are provided in Appendix A.

Under these hypotheses and definitions, the hyperelastic energy density proposed by Nolan et al.[38], and employed here in the *HGO* scenario, is given by

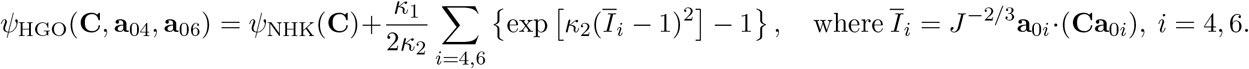

The parameters *κ*_1_ and *κ*_2_ quantify the stiffness of the fibers and their resistance to axial extension, respectively.

To allow a fair and interpretable comparison among the different constitutive models, we need connections between the rheological parameters, in particular between the linear elastic model and the neo-Hookean one. To this aim, we adopt the following expressions for the Young modulus and Poisson ratio, according to Landau et al. [39]:

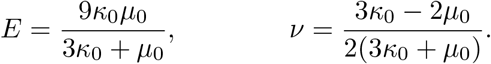

Yet, we remark that, since the parameters *κ*_0_, *µ*_0_ correspond to the volumetric and isochoric isotropic terms of the hyperelastic strain energy densities, relations (2.1) only provide a local linearization of the model.

### 2.2 Loading conditions

The reference configuration ℬ^*\**^, whose reconstruction is described in Section 3, corresponds to the condition in which the patient is lying down in the MRI scanner and is assumed to be stress-free. By this, we are implicitly neglecting the contribution of any residual stress that may reside in the tissues: this is a widely accepted assumption, motivated by both the impossibility of measuring such stress in vivo and the negligible effect it may have on macroscopic spine deformations.

Then, to compare the three computational models described above, we consider different loading conditions. Orthostatics is modeled by a uniform, downward distributed load

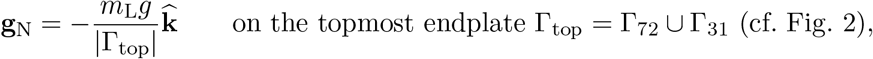

where *g* = 9.81 m s^−2^ is gravitational acceleration, *m*_L_ is the load mass, and 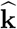 is the feet-to-head axis.

Instead, for the modeling of flexion, torsion, and lateral bending, several approaches can be found in the literature [40]. In this work, we apply pure moment conditions by means of a distributed force that depends linearly on the distance from the central vertical axis, namely

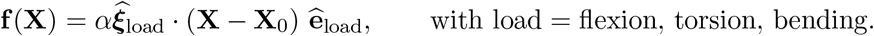

The coefficient *α* [N m^−3^] prescribes the force magnitude, **X**_0_ is the FSU barycenter while the two unit vectors 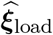, 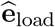 are either the right-to-left direction 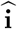, the front-to-back direction 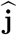, or the foot-to-head direction 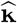, depending on the moment direction as follows: 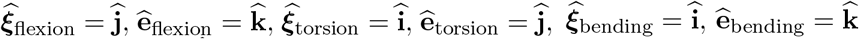, Therefore, to apply a pure momentum 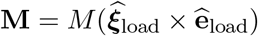we define

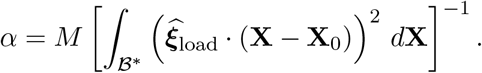

### 2.3 Numerical discretization

To solve problem (2), we employ the finite element method. We introduce the following continuous and discrete spaces

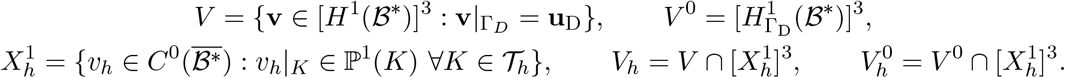

where 𝒯_*h*_ is a regular tetrahedralization of ℬ^*\**^ and ℙ^1^ denotes the space of linear polynomials [41]. Correspondingly, the discrete problem reads as follows:

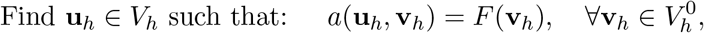

where the bilinear and linear forms are defined as

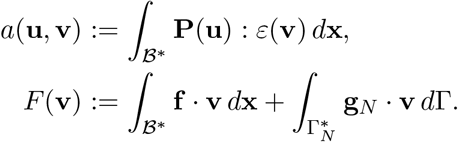

Problem (2.3) is in general nonlinear – except for the *LE* scenario, cf. Section 2.1 – therefore we employ Newton’s method to solve it. The solver is implemented in FEniCS [42], with automatic differentiation of the hyperelastic energy *ψ* by UFL’s `diff` command [43]. An absolute and relative tolerance of 10^−10^is adopted for the stopping criterion of Newton’s method, and a direct solver based on MUMPS is used to solve linear systems [44].

## 3 Geometry reconstruction and mesh generation

Volumetric lumbar MRI data of a supine patient with no lumbar pathology were obtained from the Neuroradiology Unit of Humanitas Research Hospital, in collaboration with the Department of Neurosurgery. Ethical review board approval and informed consent from the patient were obtained.

We propose a geometry reconstruction procedure that combines automatic tools and manual intervention, as detailed below, to construct a FSU encompassing two adjacent vertebrae, the annulus fibrosus of the IVD, and its nucleus pulposus. The L3 and L4 vertebrae are segmented from the images, with a resolution of 0.566 mm in all directions, whereas the soft tissue – not directly visible in MRI scans – is reconstructed in post-processing, relying on established literature data.

### 3.1 Vertebrae segmentation from MRI

The first step of the procedure is an automatic segmentation of the vertebrae using the neural network developed and trained by Levi et al. [45], consisting in a convolutional neural network with nnU-Net structure [46]. Then, the segmentation is refined using *3DSlicer* [47, 48], by local thresholds on image intensity and manual intervention on significant 2D views of the posterior processes. Finally, the refined vertebral surfaces and the images are imported in *MITK* (www.mitk.org) [49, 50], where automatic grey-level-based 3D interpolation and a smoothing filter are used to fill in the missing contours in the other image slices and remove artifacts. The resulting surfaces corresponding to vertebras L3 and L4 are exported as STL surfaces.

### 3.2 Reconstruction of intervertebral disc

For the reconstruction of the IVD we employ VMTK - Vascular Modeling ToolKit [51], with improvements developed at MOX and LaBS laboratories, Politecnico di Milano (https://github.com/checkrenzi/vmtk/tree/merge-vmtk) [52]. Starting from the vertebrae segmented in Section 3.1, we carry out the following steps:

1. first, we calculate the distance between the two adjacent vertebrae to identify and tag the inferior and superior endplates of the vertebral bodies;
2. then, we construct the outer border of the IVD connecting the tagged endplates, as an annular surface following the endplates’ edges with no bulging, since the patient is lying down in a resting position;
3. to create the inner cavity corresponding to the nucleus pulposus, we delimit an inner region on the vertebral endplates, with a relative radius of 50% [53], and we repeat the previous step to construct the annulus fibrosus inner lateral surface.

This results in a complete and conforming geometrical reconstruction of the surfaces defining the components of the FSU, which is then smoothed to remove artifacts that could hinder volumetric mesh generation. The surface tags are described in Figure 2.

**Figure 2:**
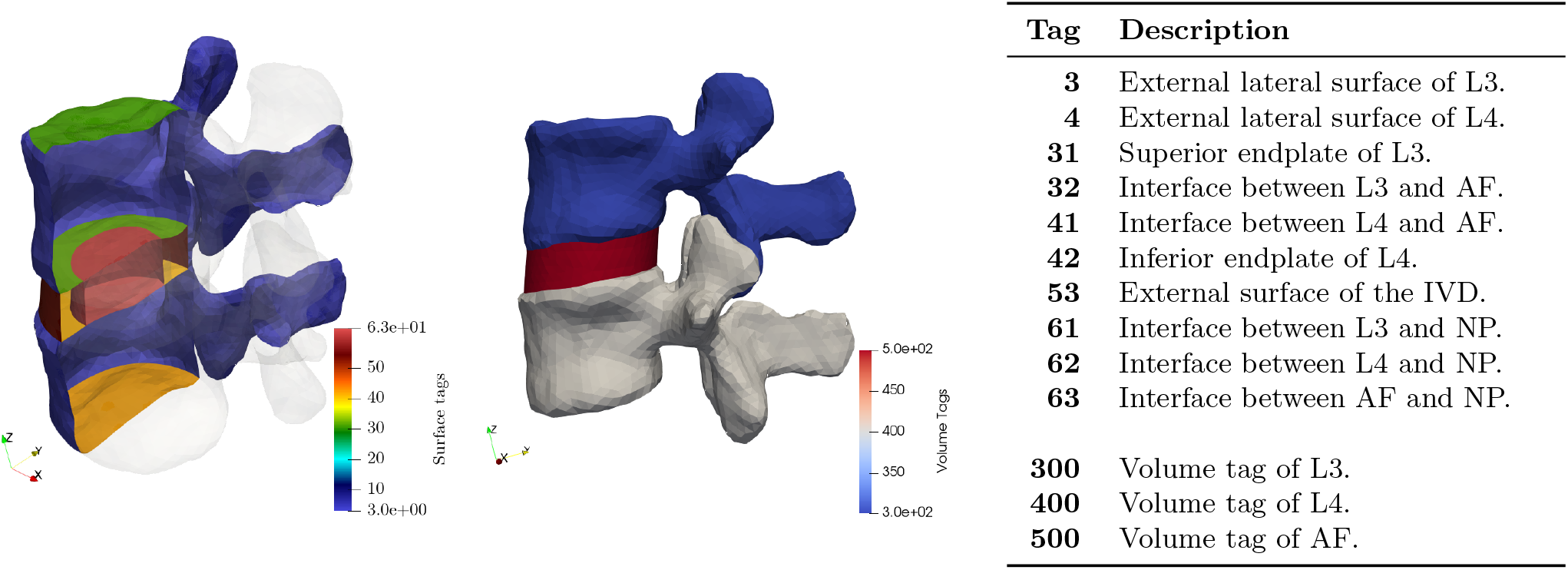
Surface tags (left) and volume tags (center) of the reconstructed FSU and their description (right).

### 3.3 Mesh generation

A tetrahedral mesh of the FSU is generated from the reconstructed surface, using VMTK. The mesh has a uniform discretization size *h* = 3 mm and it is conforming to the surface tags defined above. Different volumetric tags are defined for each vertebra and the annulus fibrosus, as described in Figure 2, to allow for different constitutive laws in the mechanical model. Fig. 3 shows the resulting mesh.

**Figure 3:**
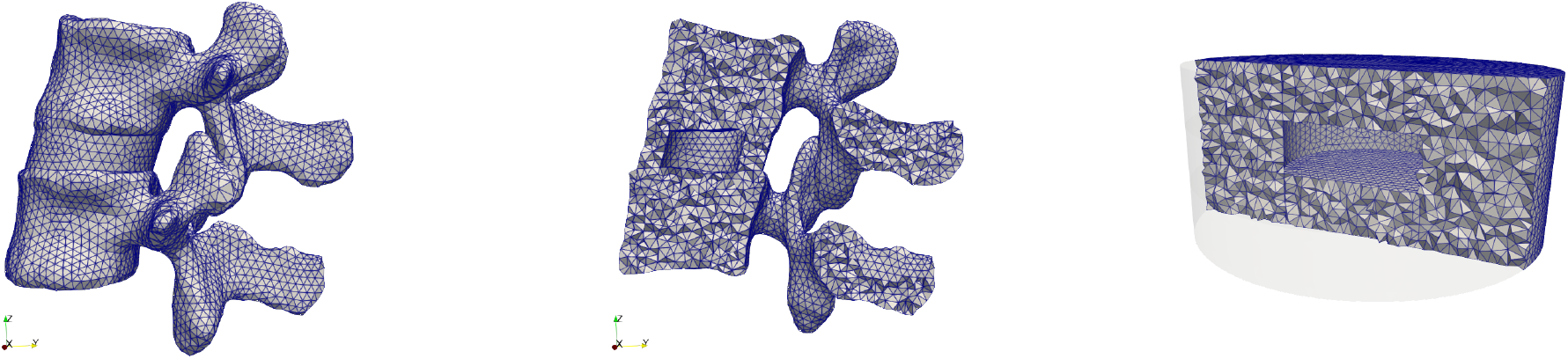
Left and center: volumetric mesh of the FSU reconstructed in Section 3 and used in the simulations of Section 5. Right: mesh of the idealized geometry used in the verification tests of Section 4, with *h* = 4.2 *×* 10^−4^ m.

## 4 Verification tests on an idealized geometry

To verify our solver, we carry out mesh convergence tests on a simplified geometry of the FSU shown in Fig. 3: three cylinders of diameter 6 cm and height 2 cm stacked along their axis, corresponding to two vertebrae and the IVD, with a cylindrical cavity of diameter 2 cm to represent the nucleus pulposus. Computational meshes of this simplified domain with different mesh sizes are generated using gmsh [54].

The tests encompass all three scenarios *LE, NH, HGO* defined in Table 2, and the mechanical parameters correspond to a physiological FSU, as reported in Table 1. We consider the following exact solution

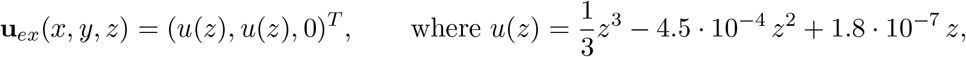

whose gradient vanishes on the vertebra-disc interfaces at *z* = 3 *·* 10^−4^ and *z* = 6 *·* 10^−4^: this is needed to prevent the discontinuities of the physical parameters from hindering the theoretically expected convergence order. Based on this solution, we compute the forcing term **f**, and we apply the following boundary conditions:

- non-homogeneous Dirichlet boundary conditions on the topmost and bottommost endplates;
- non-homogeneous Neumann boundary conditions on the remaining surfaces, namely the nucleus pulposus boundary and the external lateral surface.

The results of the convergence tests are reported in Fig. 4, and they show the theoretically expected convergence orders of 1 for the *H*^1^-error and 2 for the *L*^2^-error [41].

**Figure 4:**
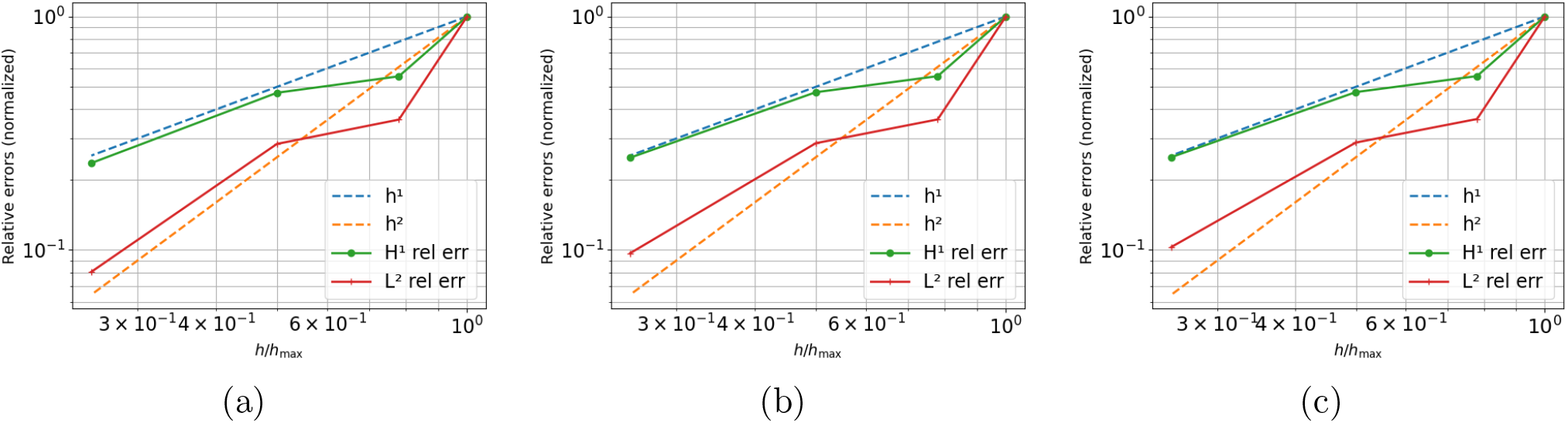
Verification tests of Section 4. Computed errors on the idealized geometry as a function of the mesh size, with *h* = (4.2, 3.3, 2.1, 1.1) *×* 10^−4^m. Referring to Table 2, the results correspond to the following scenarios: (a) *LE*, (b) *NH*, (c) *HGO*. All values are normalized with respect to the one attained for *h* = *h*_*max*_ = 4.1 *×* 10^−4^ m.

## 5 Load tests on a patient-specific lumbar L3-L4 FSU

In this section, we report and discuss the simulation of different loading conditions and assess the sensitivity of the results to significant model parameters. The methods to impose the loads has been described in Section 2.2. The comparison with literature results under flexion, bending, and torsion loads will also provide validation of the model in predicting macroscopic quantities.

### 5.1 Orthostatic parametric tests

As a first assessment test for the effect of different modeling choices, we report the simulation of orthostatic loading, with different values of the load mass *m*_L_ = 20, 40, 80 kg. No volumetric load is applied (**f** = **0**), the bottommost endplate Γ_bot_ is kept fixed (**u** = **0**) and the remaining boundaries are traction-free (**P**(**u**)**n** = **0**).

In Fig. 5, we report the displacement field and deformed configuration under such orthostatic loads for each of the three scenarios of Table 2. In all cases, the strongest deformation is observed in the spinous process of L3, whereas the whole L4 is essentially undeformed (maximum deformation over L4 < 0.04 mm), while being fully constrained only at its bottom endplate. Moreover, the distribution of the deformation field essentially corresponds to a backward flexion in response to orthostatic loads. This is consistent with clinical observations and the fact that the most compliant portion of the FSU is the IVD. Indeed, for each loading condition, we can observe that the maximum deformation magnitude is attained in the *LE* scenario (max **u**_*LE*_ = 1.52 mm for *m*_L_ = 20 kg), which has the most compliant AF: the nonlinear hyperelastic rheology of the *NH* and *HGO* models yields a significant stiffening of the AF (max **u**_*NH*_ = 1.44 mm, max **u**_*HGO*_ = 1.43 mm for *m*_L_ = 20 kg).

**Figure 5:**
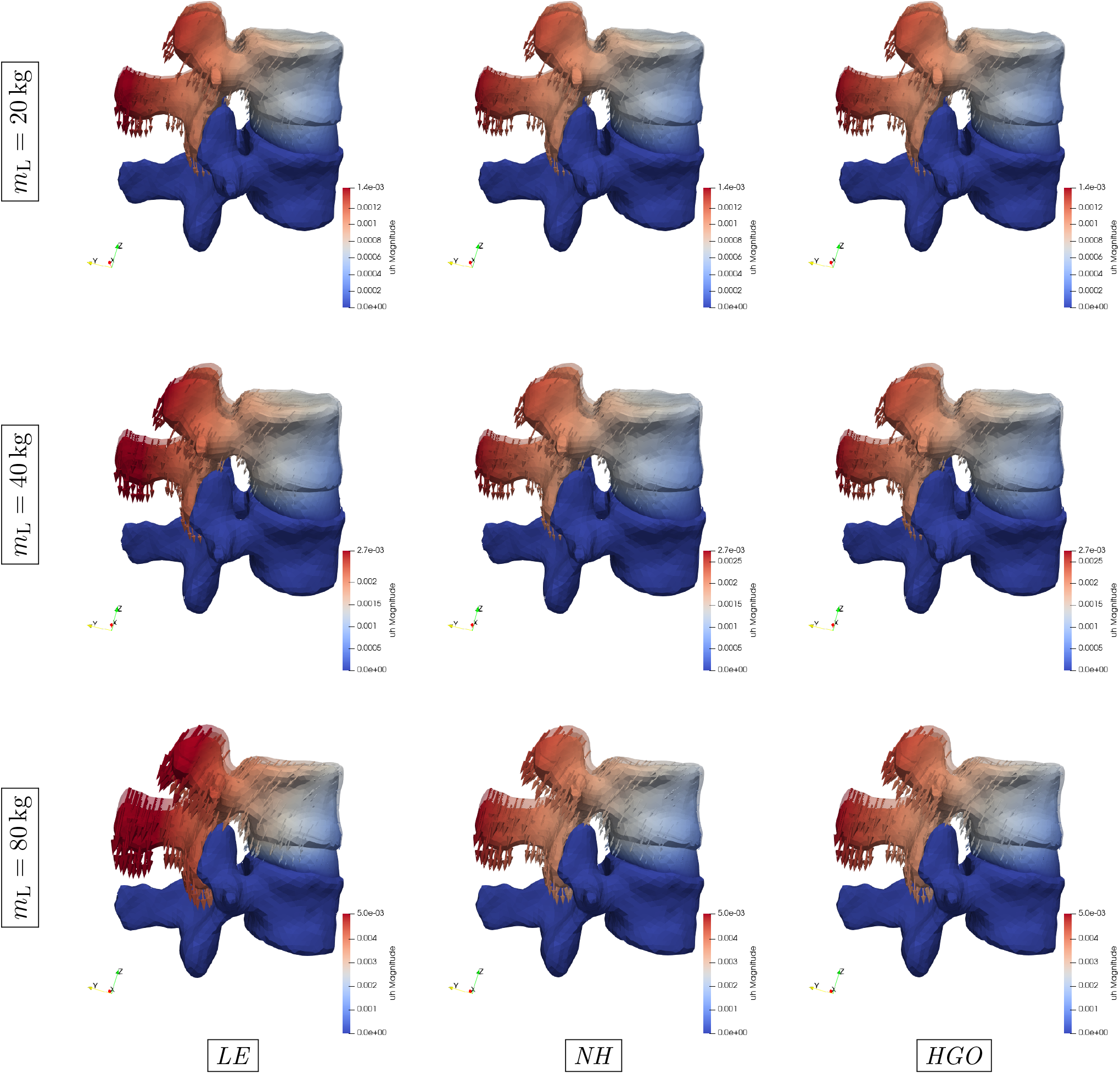
Deformed configuration and displacement field in the orthostatic tests of Section 5.1 for the three scenarios of Table 2 (from left to right), under a load of *m*_L_ = 20, 40, 80 kg (from top to bottom). The reference, unloaded configuration is in transparency. To better display the differences among the three scenarios, the arrows are exaggerated by a factor of 7 (top, *m*_L_ = 20 kg), 4 (middle, *m*_L_ = 40 kg), 3 (bottom, *m*_L_ = 80 kg).

In terms of the effects of the fiber structure of the *HGO*, we point out that only a slight effect seems to be visible, in terms of L3 displacement: as shown in Fig. 6 for the case with the largest load mass *m*_L_ = 80 kg, the differences between *LE* and *NH* are much larger than those between the two hyperelastic models. However, focusing on the AF, we can notice that fiber reinforcement increases the resistance to bulging: as displayed in Fig. 7, the *HGO* scenario shows significantly less AF bulging than the other two, which is particularly evident in the coronal view. This is the main reason that is given in the literature to adopt fiber-reinforced constitutive models [9, 13, 40, 55, 18].

**Figure 6:**
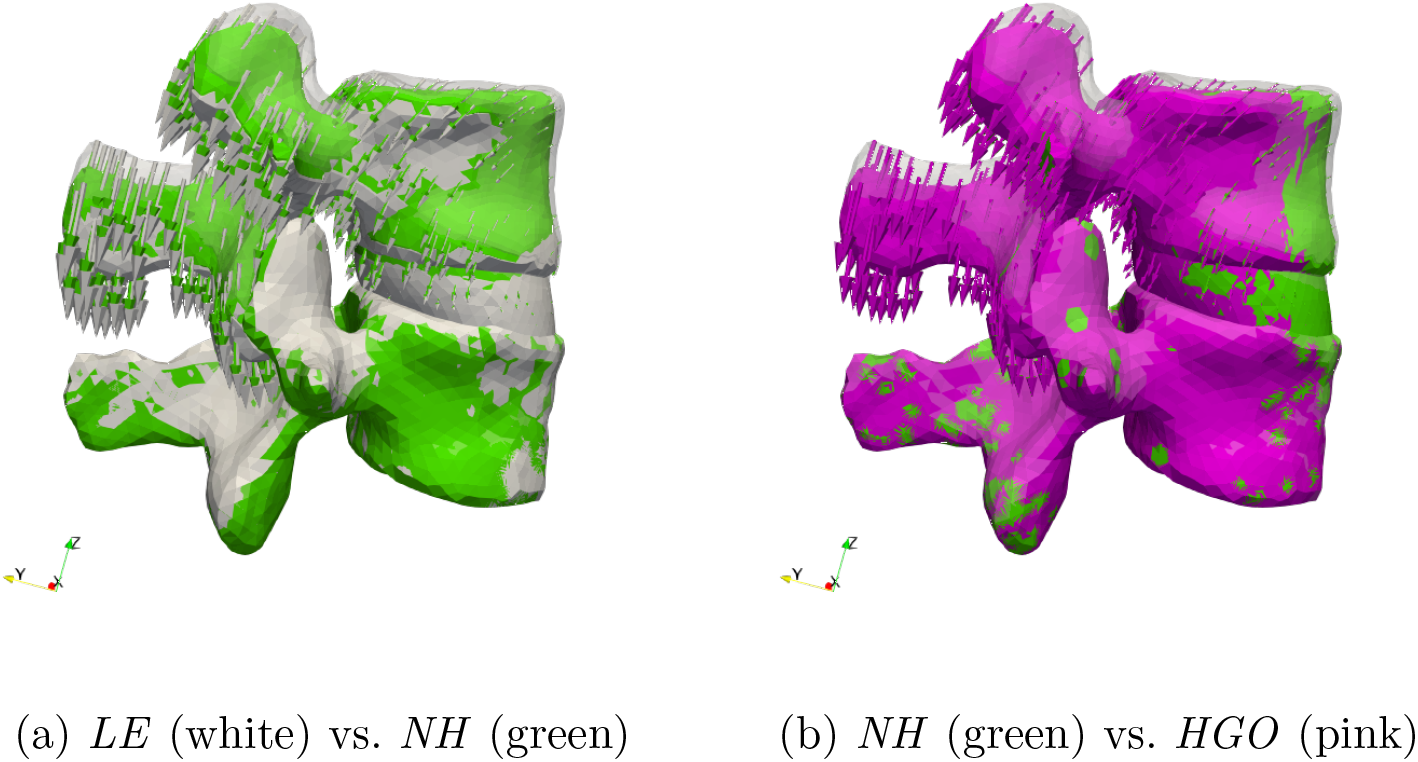
Pair-wise comparison of mechanical models under the orthostatic load of Section 5.1 with *m*_L_ = 80 kg. The deformed configurations are represented in solid colors, while the reference configuration is in transparency.

**Figure 7:**
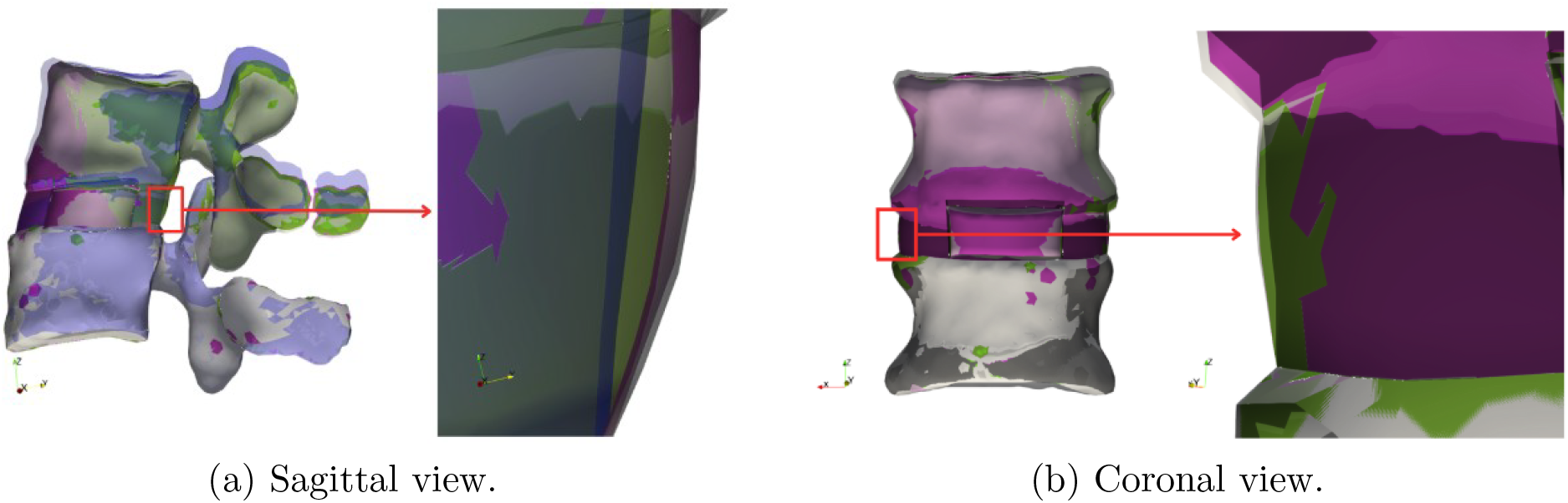
Comparison between the constitutive models under an orthostatic load of *m*_L_ = 80 kg. The reference configuration is in blue (in (a)) and black (in (b)), while the white, green, and pink correspond to the deformed configuration of *LE, NH*, and *HGO*, respectively.

To better quantify the differences among the models, we assess the overall shortening of the FSU in terms of the average vertical displacement of the topmost endplate Γ_top_, defined as

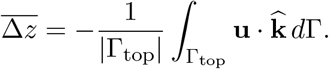

As shown in Fig. 8, we can observe a substantially linear dependence of 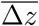 from *m*_L_. The slope of this curve is slightly reduced in the case of the hyperelastic models incorporating the nonlinear and anisotropic properties of the AF, due to the aforementioned disc stiffening effect. This is consistent with the differences among the model being more evident for larger values of the loading mass.

**Figure 8:**
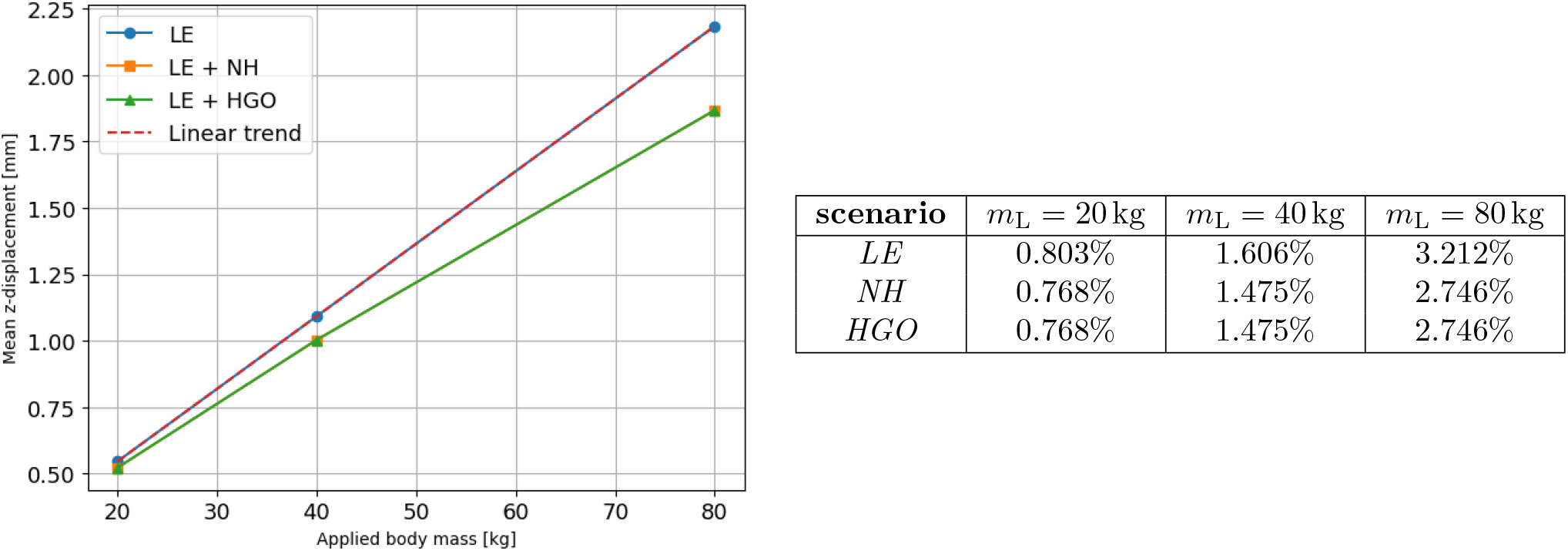
Orthostatic tests of Section 5.1. Average vertical shortening 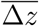 in the three scenarios, in absolute value (left figure) and as a percentage of the initial overall axial length of the FSU (right table), as a function of the loading mass *m*_L_. In the plot, the lines of *NH* and *HGO* are overlapping.

### 5.2 Realistic loading and validation

In this section, we go beyond orthostatics and present numerical experiments in several active deformation settings. In particular, Section 5.2.1 addresses a pure forward flexion around the coronal axis, while lateral bending and pure torsion are discussed in sections 5.2.2 and 5.2.3, respectively. As mentioned in Section 2.2, the specific loading conditions are achieved by a distributed volumetric force **f** applied only to L3 (see Fig. 9), while **f** = **0** in L4. The bottom endplate of L4 is fixed, the top endplate of L3 is loaded with an orthostatic mass of *m*_L_ = 80 kg, and the remaining boundaries are traction-free.

**Figure 9:**
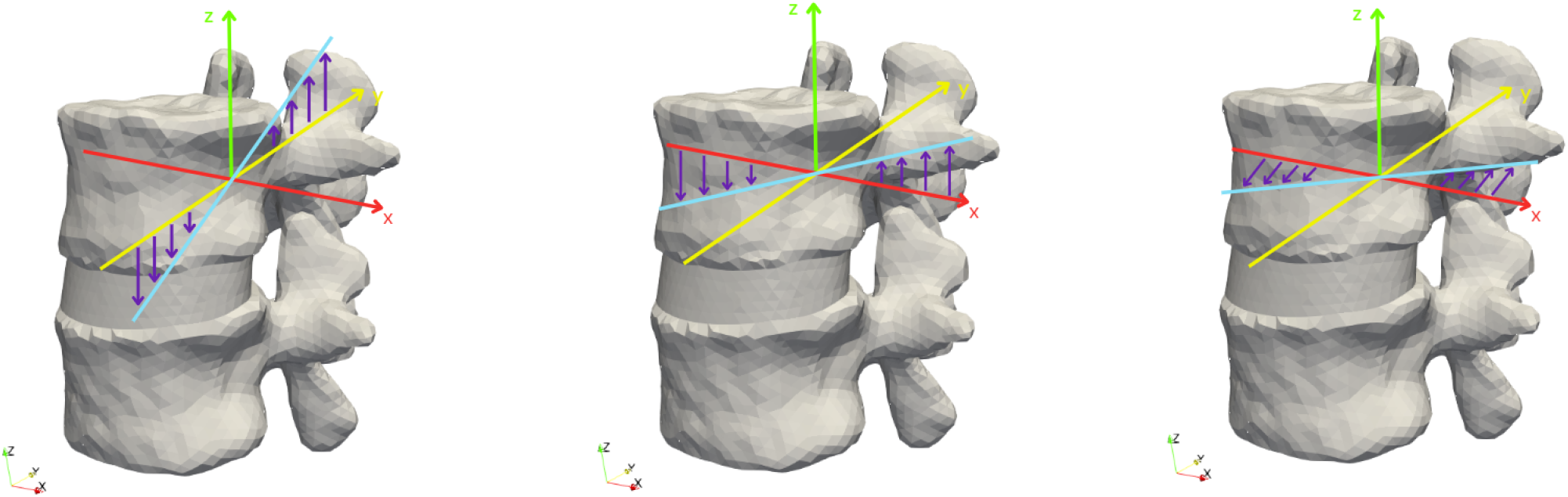
Graphical representation of the applied force *f* in the case of pure flexion (left – Section 5.2.1), lateral bending (center – Section 5.2.2), and pure torsion (right – Section 5.2.3).

In each of the following tests, we compare the results obtained in the three scenarios *LE, NH, HGO*, and we also discuss a macroscopic validation against literature results. To this aim, in addition to present and discuss the distribution of the displacement field, we also analyze the Range Of Motion (ROM), which is used to quantify the overall movement of the FSU[13]. This is defined as the plane angle between the axis of L3 in the deformed configuration and the one in the reference configuration – the axis being the line connecting the two centroids of the endplates of L3.

#### 5.2.1 Flexion

In this section, we consider a loading state combining the orthostatic load with *m*_L_ = 80 kg as in Section 5.1 and a pure forward flexion around the coronal axis as depicted in Fig. 9. We first report a parametric analysis with respect to the flexion moment magnitude, just for the *HGO* scenario, and then compare the results of the three scenarios *LE, NH, HGO*, with literature results to make a step towards validation.

For the parametric study, three different flexion moments are applied to the FSU, with *M* = 4, 7.5, 11 N m (cf. (2.2)), obtaining the results reported in Fig. 10. We can observe that a moment of small magnitude (*M* = 4 N m) simply counteracts the small backward flexion component of the orthostatic load observed in Fig. 5, resulting in a similar condition to pure axial compression. Stronger moments, instead, dominate the orthostatic effects and show a clear forward flexion. To quantify this effect, we compute the ROM in the three cases, as displayed in Fig. 11: the resulting angles show a linear dependence of the ROM on the moment magnitude *M*.

**Figure 10:**
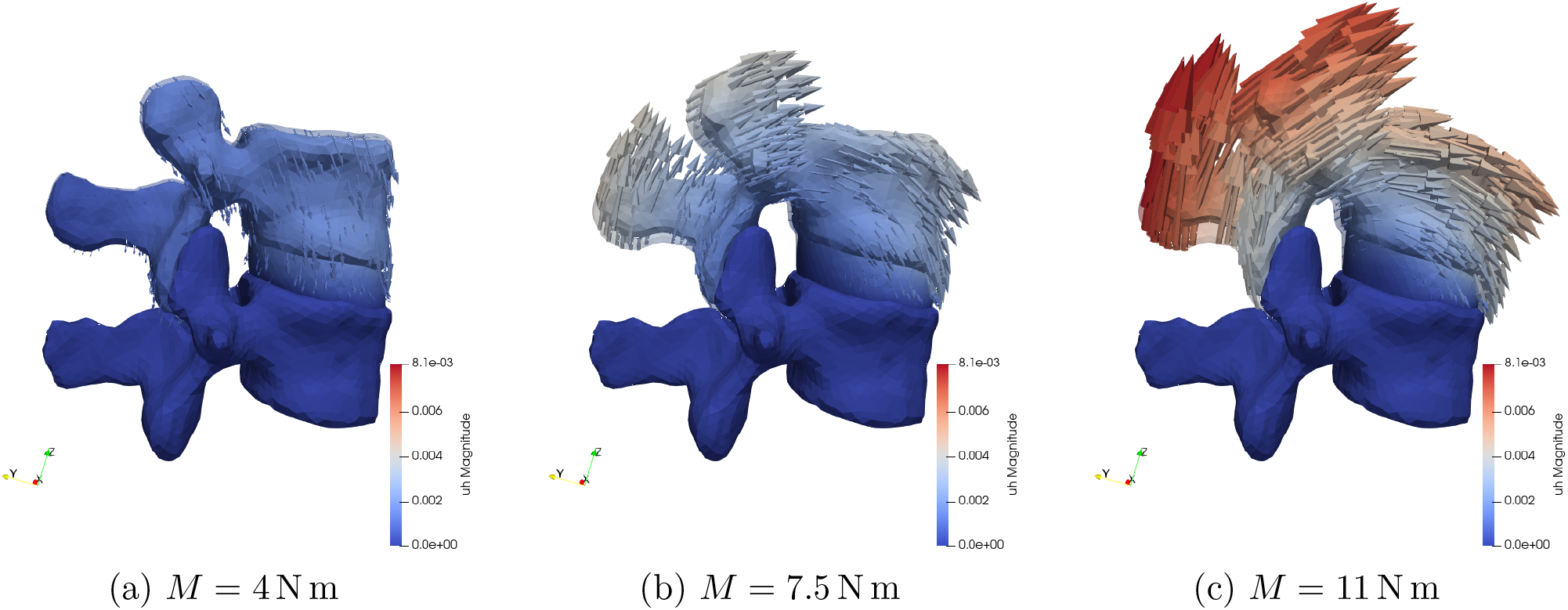
Pure flexion test of Section 5.2.1 – *HGO* scenario. Deformed configuration and displacement field. The reference, unloaded configuration is in transparency, and the arrows are exaggerated by a factor 3.

**Figure 11:**
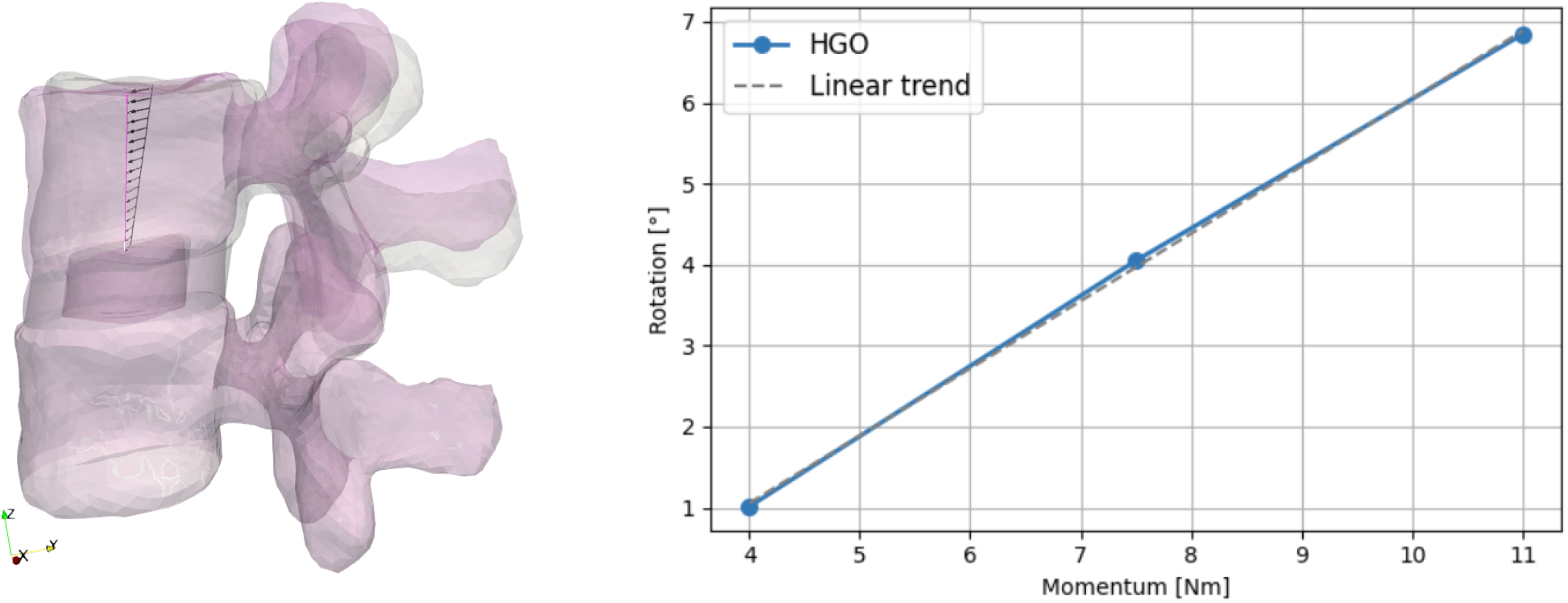
Pure flexion test of Section 5.2.1 – *HGO* scenario. Left: graphical representation of the Range Of Motion (ROM) highlighted between the reference (in white) and the deformed (in pink) configuration, for *M* = 7.5 N m. Right: ROM trend with respect to increasing momentum *M*.

As a validation step, we make a comparison between our results in the different scenarios and those reported by Rohlmann et al. [40], with the same flexion moment *M* = 7.5 N m applied in that study. The ROM values under this load are the following:

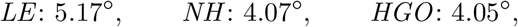

which are all in the range of 4.0° − 6.0° reported by Rohlmann et al. [40]. As in the orthostatic tests above, we observe that the *LE* model is more compliant than the nonlinear hyperelastic ones, yet all ROM values are consistent with the literature.

#### 5.2.2 Lateral bending

The loading state of this section is a pure lateral bending moment along the antero-posterior axis (see (2.2) and Fig. 9, center), together with an orthostatic load of *m*_L_ = 80 kg, and we start focusing on the *HGO* scenario. To allow comparison with literature data, we apply a moment of 7.8 N m as employed by Dreischarf et al. [56]. In Fig. 12, we report the resulting displacement, showing an expected bending to the right, with a stronger magnitude of the displacement on the compressed side, with respect to the stretched side. This is consistent with the role of fiber reinforcement in the *HGO* constitutive relation, which makes the AF stiffer under stretching strains than under compression [28].

**Figure 12:**
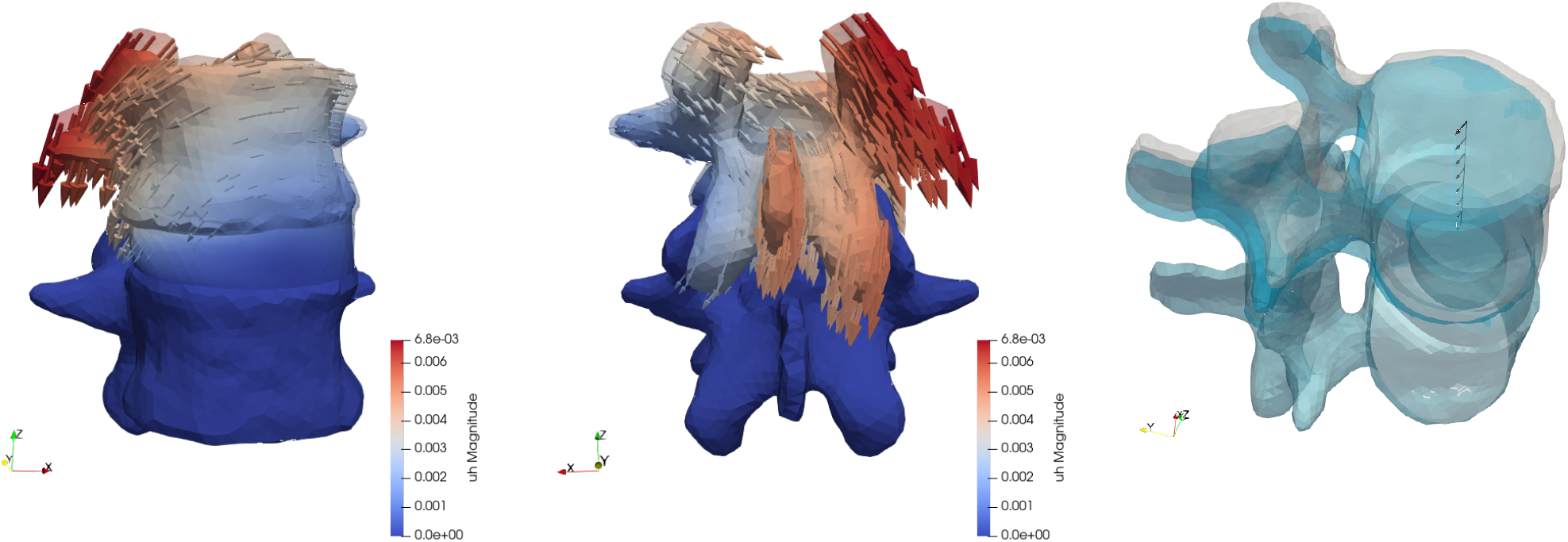
Lateral bending test of Section 5.2.2 – *HGO* scenario. Deformed configuration and displacement field. The reference, unloaded configuration is in transparency, and the arrows are exaggerated by a factor 3. On the right, the ROM shows combined contributions of bending, backward flexion, and slight torsion, as highlighted by the black arrows.

Computing the ROM in this case, as displayed in Fig. 12-right, we obtain an angle of 5.28°, which is very close to the average value of the in-vivo reference values range of [4.3°, 6.3° reported by Dreischarf et al. [56]. We point out that this rotation is not strictly confined to the coronal plane, but it is rather a combination of lateral bending (−4.69° in the coronal plane XZ) and backward flexion (−2.93° in the sagittal plane YZ). Moreover, a secondary effect of this combined load is also a slight torsion of L3 about the vertical axis, which is commonly noted in in-vivo clinical observations.

Finally. to investigate the effect of the modeling choices, we compare apply the same loading conditions also in the *LE* and *NH* scenarios, obtaining the following ROM values:

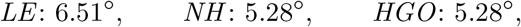

As in the previous sections, we observe significantly more compliance in the *LE* scenario, which exceeds the span of physiological values reported by Dreischarf et al. [56]. This comparison further supports the adoption of a nonlinear hyperelastic model for the AF.

#### 5.2.3 Torsion

As a final test, we apply a torsion moment of *M* = 5.5 N m to L3, together with an orthostatic load of *m*_L_ = 80 kg, in the *HGO* scenario. The resulting displacement field, reported in Fig. 13, is stronger than all previous tests, reaching a maximum magnitude of 1.2 cm in the spinous process. We notice an asymmetric deformation pattern, which is physically consistent with the superposition of orthostatics and torsion: on the left side, the purely torsional rotation has a positive antero-posterior component which adds up to the orthostatic-induced backward flexion, while on the right side this component is directed forward, thus partially opposing such flexion.

**Figure 13:**
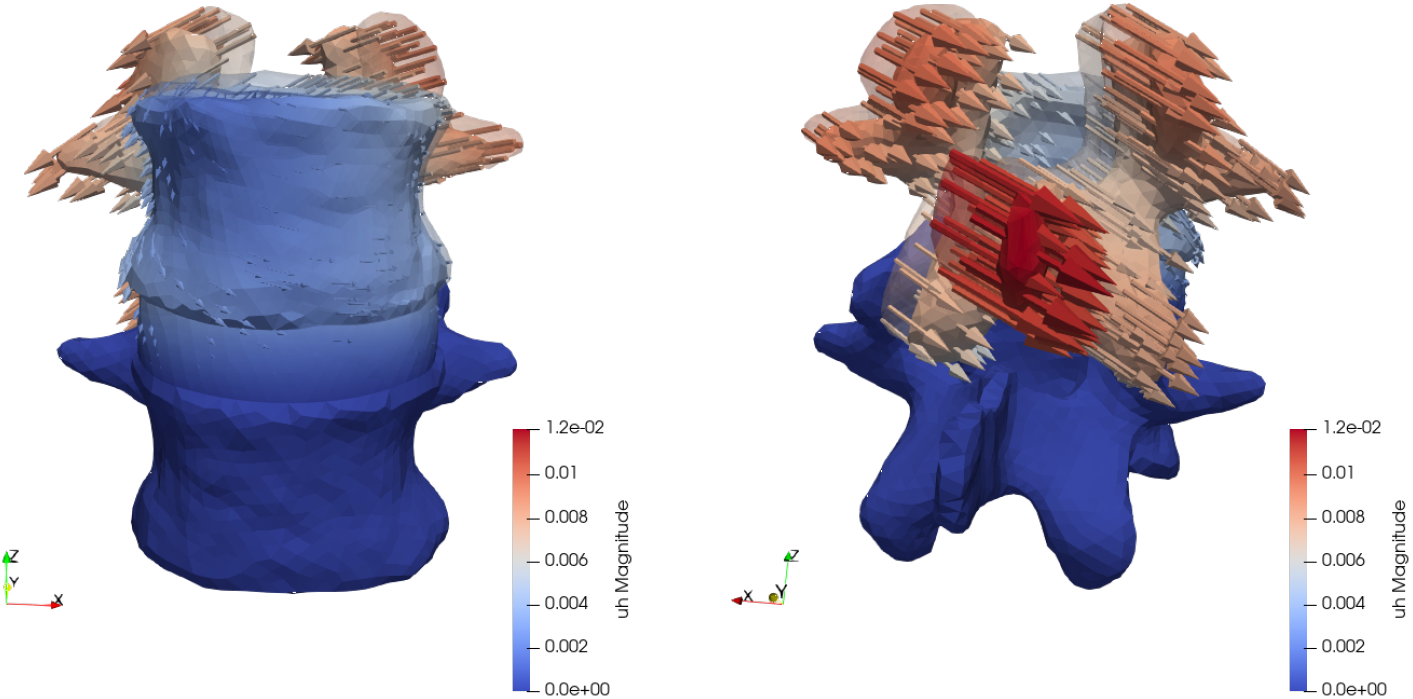
Torsion test of Section 5.2.3 – *HGO* scenario. Deformed configuration and displacement field. The reference, unloaded configuration is in transparency, and the arrows are exaggerated by a factor 2.

In terms of angles, this relatively large displacement is mostly torsional, as expected: L3 undergoes a rotation about the vertical axis of 7.7°, while the ROM – displayed in Fig. 14 is just 3.89°. The latter shows that a purely torsional load also induces non-negligible flexion and lateral bending of the FSU, which is consistent with what is commonly observed in computational and in-vivo assessments, yet these secondary effects are less than those obtained in the actual flexion and lateral bending tests of Sections 5.2.1 and 5.2.2. Comparing the results with the literature, we observe that our model significantly overestimates the torsion angle of 1.1° reported by Dreischarf et al. [57]. This discrepancy could be ascribed to the lack of facet joints in our model, whose main functional role is indeed to stabilize against intervertebral relative rotation.

**Figure 14:**
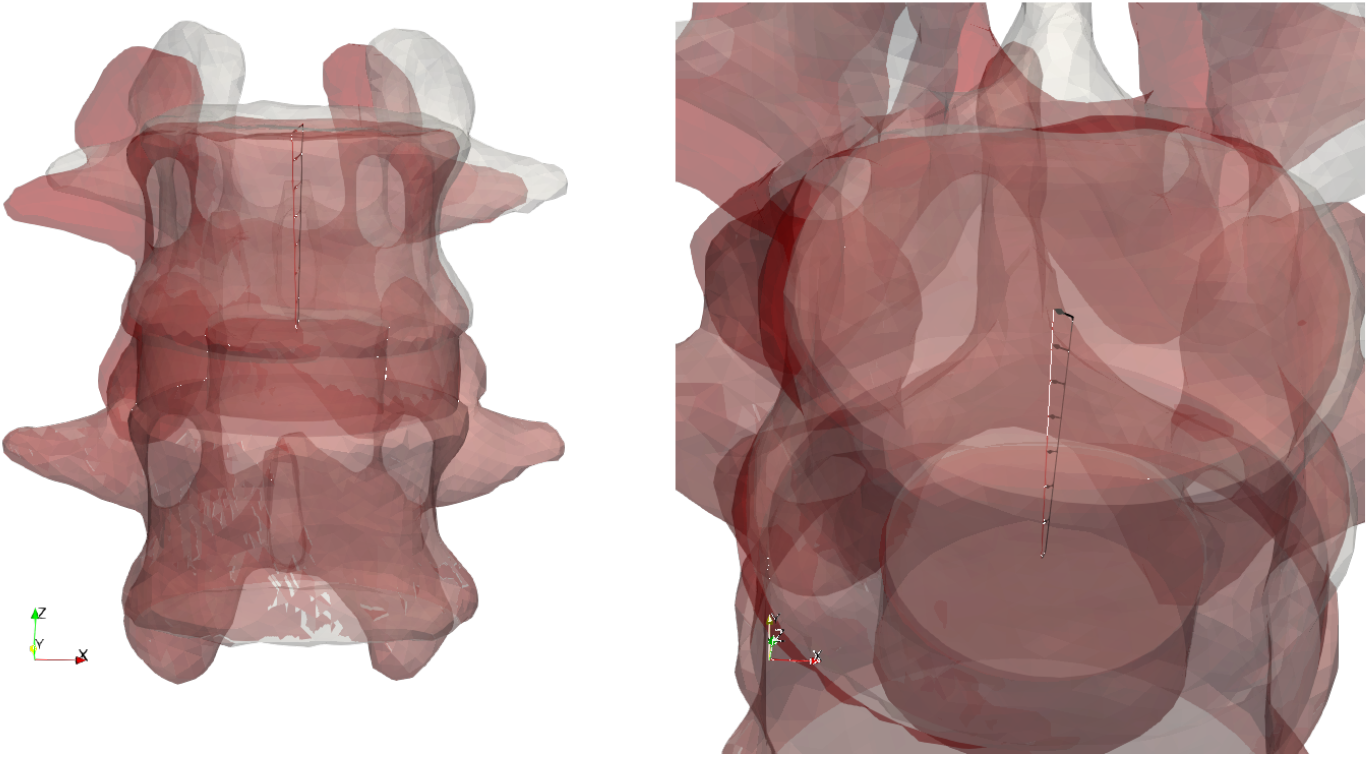
Torsion test of Section 5.2.3 – *HGO* scenario. Graphical representation of the Range of motion (ROM) highlighted between the reference (in white) and the deformed (in red) configuration.

## 6 Conclusions

In this work, we presented the development, verification, and preliminary validation of a finite element model for the study of spinal biomechanics on patient-specific geometries. Verification tests on an idealized geometry assessed and confirmed the accuracy of the computational model. Starting from clinical MRI images, a detailed segmentation of the vertebral bodies and the reconstruction of the intervertebral disc was carried out, yielding a realistic three-dimensional model of the L3-L4 functional spinal unit (FSU). In this FSU, the vertebrae’s mechanical response was modeled by a linear elastic relation, while three constitutive models for the Annulus Fibrosus (AF) were compared: linear elasticity (*LE*), Neo-Hookean isotropic hyperelasticity (*NH*), and a fiber-reinforced Holzapfel-Gasser-Ogden model (*HGO*), calibrated with material parameters drawn from the literature. [9]

The employment of a real, reconstructed geometry allowed to assess the mechanical response of the different components of the FSU with anatomical accuracy. The simulations performed under orthostatic loading, pure flexion, and lateral bending, reproduced physiologically meaningful deformation patterns and Range-Of-Motion angles in good agreement with reference values reported in the literature. [40, 57] Moreover, they quantitatively assessed the impact of the AF’s constitutive model on the macroscopic mechanical response of the FSU: the linear constitutive relation yields an overestimation of the FSU compliance, therefore a nonlinear model such as *NH* or *HGO* is required to reproduce physiological results.

An additional test under torsion loading provided physically sound results but highlighted the limitation of not considering facet joints in the model. The extension of the model to include facet joints – as well as other ligaments – in an anatomically accurate way requires significant enhancement of the geometry reconstruction pipeline and a suitable local refinement of the computational mesh to account for the small thickness of ligaments, and is thus postponed to future work. Another extension that could improve the model’s ability to reproduce physiological response is the introduction of a compliance model for the nucleus pulposus, to better account for pressure redistribution in the IVD. Furthermore, more refined boundary conditions – e.g. including the follower load from other vertebrae [56], or accounting for lateral loads related to muscle activation – would allow to reproduce several loading conditions both in physiological and pathological conditions. Such boundary conditions could also act as a surrogate of portions of the spinal column that are not directly included in the reconstructed geometry, thus keeping under control the anatomical reconstruction step, which is the most time consuming part of the computational pipeline. Finally, all these extensions would further enhance the predictive capability of the model, making it better suitable to support diagnosis and clinical decisions.

## Acknowledgments

IF and PFA have been partially supported by ICSC-Centro Nazionale di Ricerca in High Performance Computing, Big Data, and Quantum Computing funded by European Union–NextGenerationEU (CUP D43C22001240001). The present research is part of the activities of “Dipartimento di Eccellenza 2023-2027”, Dipartimento di Matematica, Politecnico di Milano. IF, SP, and PFA are members of GNCS-INdAM.

## A Fiber orientation

In this section, we detail the procedure employed to define fiber orientation in the annulus fibrosus. As reported in the literature [9], the fibers lie orthogonally to the radial direction **ê**_*r*_ and have an angle with respect to the vertical direction **ê** _*z*_ that progressively passes from 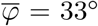 on the outer surface to 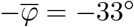 on the internal interface with the nucleus pulposus.

If we model the annulus as perfectly cylindrical, with internal radius *R*_1_ and external radius *R*_2_, the fiber direction can be identified by the unit vector

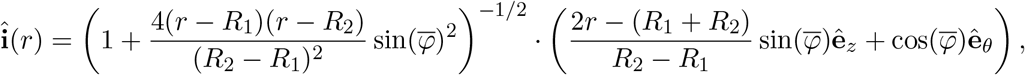

where *r* is the radial coordinate and **ê**_*θ*_ = **ê**_*z*_ *×* **ê**_*r*_. The resulting vector field is displayed in Fig. 15.

**Figure 15:**
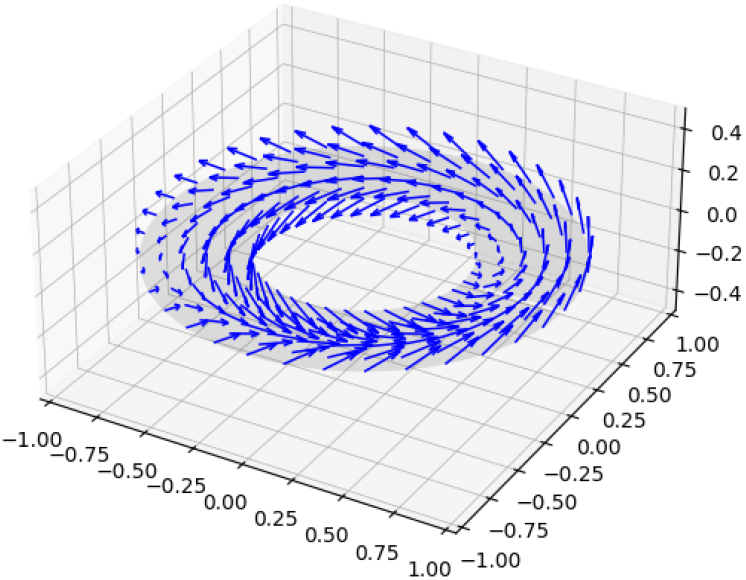
Fibers’ orientation in idealized geometry.

In cartesian coordinates, the fiber orientation vector reads as follows:

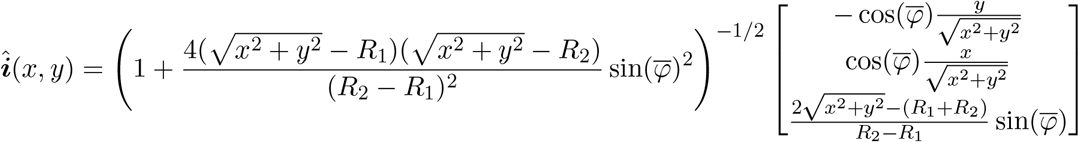

To employ this fiber distribution in the real disc geometry reconstructed in Section 3.2, we consider the disc centroid as the origin, align the *z*-axis to the axis of the disc, and set the parameters *R*_1_ and *R*_2_ to the *average* values of the outer and inner radius, respectively.

## References

[1] M. L. Ferreira, K. De Luca, L. M. Haile, J. D. Steinmetz, G. T. Culbreth, M. Cross, J. A. Kopec, P. H. Ferreira, F. M. Blyth, R. Buchbinder, et al., “Global, regional, and national burden of low back pain, 1990–2020, its attributable risk factors, and projections to 2050: a systematic analysis of the global burden of disease study 2021,” The Lancet Rheumatology, vol. 5, no. 6, pp. e316–e329, 2023.

[2] M. Cheng, Y. Xue, M. Cui, X. Zeng, C. Yang, F. Ding, and L. Xie, “Global, regional, and national burden of low back pain: findings from the global burden of disease study 2021 and projections to 2050,” Spine, vol. 50, no. 7, pp. E128–E139, 2025.

[3] I. Urits, A. Burshtein, M. Sharma, L. Testa, P. A. Gold, V. Orhurhu, O. Viswanath, M. R. Jones, M. A. Sidransky, B. Spektor, et al., “Low back pain, a comprehensive review: pathophysiology, diagnosis, and treatment,” Current pain and headache reports, vol. 23, no. 3, p. 23, 2019.

[4] G. K. Harada, Z. K. Siyaji, S. Younis, P. K. Louie, D. Samartzis, and H. S. An, “Imaging in spine surgery: Current concepts and future directions,” Spine Surgery and Related Research, vol. 4, no. 2, pp. 99–110, 2020.

[5] G. Capo, F. Calvanese, N. Tahhan, D. Creatura, I. Zaed, E. Bellina, A. Baram, F. Cotton, and C. Y. Barrey, “Prediction of MRI in intra-operative findings for spinal meningeal diseases,” Neurochirurgie, vol. 71, no. 3, p. 101661, 2025.

[6] A. E. Levent, M. Tanaka, C. Kumawat, C. Heng, S. Nikolaos, K. Latka, A. Miyamoto, T. Komatsubara, S. Arataki, Y. Oda, K. Shinohara, and K. Uotani, “Review article: Diagnostic paradigm shift in spine surgery,” Diagnostics, vol. 15, no. 5, 2025.

[7] A. G. Bruno, M. L. Bouxsein, and D. E. Anderson, “Development and validation of a musculoskeletal model of the fully articulated thoracolumbar spine and rib cage,” Journal of biomechanical engineering, vol. 137, no. 8, p. 081003, 2015.

[8] D. Ignasiak, S. Dendorfer, and S. J. Ferguson, “Thoracolumbar spine model with articulated ribcage for the prediction of dynamic spinal loading,” Journal of biomechanics, vol. 49, no. 6, pp. 959–966, 2016.

[9] E. Bellina, M. E. Laurino, A. Perego, A. Pezzinga, L. Carpenedo, D. Ninarello, and L. La Barbera, “Assessment of a fully-parametric thoraco-lumbar spine model generator with articulated ribcage,” Journal of Biomechanics, vol. 164, p. 111951, 2024.

[10] Y. Su, A. I. Tsirikos, V. Koutsos, and P. Pankaj, “Development and assessment of a novel generic finite element spine model for clinical applications,” International Journal for Numerical Methods in Biomedical Engineering, vol. 41, no. 9, p. e70098, 2025.

[11] M. Iwamoto, Y. Nakahira, and H. Kimpara, “Development and validation of the total human model for safety (thums) toward further understanding of occupant injury mechanisms in precrash and during crash,” Traffic injury prevention, vol. 16, no. sup1, pp. S36–S48, 2015.

[12] I. E. Bojairami, K. El-Monajjed, and M. Driscoll, “Development and validation of a timely and representative finite element human spine model for biomechanical simulations,” Scientific Reports, vol. 10, p. 21519, 2020.

[13] M. Dreischarf, T. Zander, A. Shirazi-Adl, C. Puttlitz, C. Adam, C. Chen, V. Goel, A. Kiapour, Y. Kim, K. Labus, J. Little, W. Park, Y. Wang, H. Wilke, A. Rohlmann, and H. Schmidt, “Comparison of eight published static finite element models of the intact lumbar spine: Predictive power of models improves when combined together,” Journal of Biomechanics, vol. 47, no. 8, pp. 1757–1766, 2014.

[14] D. J. Coombs, P. J. Rullkoetter, and P. J. Laz, “Efficient probabilistic finite element analysis of a lumbar motion segment,” Journal of Biomechanics, vol. 61, pp. 65–74, 2017.

[15] A. Rohlmann, A. Mann, T. Zander, et al., “Effect of an artificial disc on lumbar spine biomechanics: a probabilistic finite element study,” European Spine Journal, vol. 18, pp. 89–97, 2009.

[16] W. Wang, C. Zhou, R. Guo, T. Cha, and G. Li, “Prediction of biomechanical responses of human lumbar discs - a stochastic finite element model analysis,” Computer Methods in Biomechanics and Biomedical Engineering, vol. 24, no. 15, pp. 1730–1741, 2021. PMID: 34121532.

[17] M. Y. Lu, W. C. Hutton, and V. M. Gharpuray, “Can variations in intervertebral disc height affect the mechanical function of the disc?,” Spine, vol. 21, no. 19, pp. 2208–2216, 1996.

[18] U. M. Ayturk and C. M. Puttlitz, “Parametric convergence sensitivity and validation of a finite element model of the human lumbar spine,” Computer methods in biomechanics and biomedical engineering, vol. 14, no. 8, pp. 695–705, 2011.

[19] C. Du, Z. Mo, S. Tian, L. Wang, J. Fan, S. Liu, and Y. Fan, “Biomechanical investigation of thoracolumbar spine in different postures during ejection using a combined finite element and multi-body approach,” International journal for numerical methods in biomedical engineering, vol. 30, no. 11, pp. 1121–1131, 2014.

[20] C. Garavelli, C. Curreli, M. Palanca, A. Aldieri, L. Cristofolini, and M. Viceconti, “Experimental validation of a subject-specific finite element model of lumbar spine segment using digital image correlation,” PLOS ONE, vol. 17, pp. 1–17, 09 2022.

[21] C.-K. Lee, Y. E. Kim, C.-S. Lee, Y.-M. Hong, J.-M. Jung, and V. K. Goel, “Impact response of the intervertebral disc in a finite-element model,” Spine, vol. 25, no. 19, pp. 2431–2439, 2000.

[22] M. Rasouligandomani, A. del Arco, F. K. Chemorion, et al., “Dataset of finite element models of normal and deformed thoracolumbar spine,” Scientific Data, vol. 11, p. 549, 2024.

[23] G. A. Holzapfel, C. A. Schulze-Bauer, G. Feigl, and P. Regitnig, “Single lamellar mechanics of the human lumbar anulus fibrosus,” Biomechanics and modeling in mechanobiology, vol. 3, no. 3, pp. 125–140, 2005.

[24] H. Schmidt, F. Galbusera, A. Rohlmann, T. Zander, and H.-J. Wilke, “Effect of multilevel lumbar disc arthroplasty on spine kinematics and facet joint loads in flexion and extension: a finite element analysis,” European spine journal, vol. 21, no. Suppl 5, pp. 663–674, 2012.

[25] T. Wiczenbach, L. Pachocki, K. Daszkiewicz, P. L uczkiewicz, and W. Witkowski, “Development and validation of lumbar spine finite element model,” PeerJ, vol. 11, p. e15805, 2023.

[26] R. Wang and Z. Wu, “Recent advancement in finite element analysis of spinal interbody cages: A review,” Frontiers in Bioengineering and Biotechnology, vol. Volume 11 - 2023, 2023.

[27] R. Eberlein, G. A. Holzapfel, and C. A. Schulze-Bauer, “An anisotropic model for annulus tissue and enhanced finite element analyses of intact lumbar disc bodies,” Computer methods in biomechanics and biomedical engineering, vol. 4, no. 3, pp. 209–229, 2001.

[28] T. C. Gasser, R. W. Ogden, and G. A. Holzapfel, “Hyperelastic modelling of arterial layers with distributed collagen fibre orientations,” Journal of the Royal Society Interface, vol. 3, no. 6, pp. 15–35, 2006.

[29] J. Campbell, D. Coombs, M. Rao, P. Rullkoetter, and A. Petrella, “Automated finite element meshing of the lumbar spine: Verification and validation with 18 specimen-specific models,” Journal of Biomechanics, vol. 49, no. 13, pp. 2669–2676, 2016.

[30] S. Caprara, F. Carrillo, J. G. Snedeker, M. Farshad, and M. Senteler, “Automated pipeline to generate anatomically accurate patient-specific biomechanical models of healthy and pathological FSUs,” Frontiers in Bioengineering and Biotechnology, vol. Volume 9 - 2021, 2021.

[31] E. Biamonte, R. Levi, F. Carrone, et al., “Artificial intelligence-based radiomics on computed tomography of lumbar spine in subjects with fragility vertebral fractures,” Journal of Endocrinological Investigation, vol. 45, pp. 2007–2017, 2022. Version of record: 25 June 2022. Issue date: October 2022.

[32] X. He, Y. Qiu, X. Lai, Z. Li, L. Shu, W. Sun, and X. Song, “Towards a shape-performance integrated digital twin for lumbar spine analysis,” Digital Twin, vol. 2, no. 2, p. 2520116, 2025.

[33] M. Ahmadi, D. Biswas, R. Paul, M. Lin, Y. Tang, T. S. Cheema, E. D. Engeberg, J. Hashemi, and F. D. Vrionis, “Integrating finite element analysis and physics-informed neural networks for biomechanical modeling of the human lumbar spine,” North American Spine Society Journal (NASSJ), vol. 22, p. 100598, June 2025.

[34] J. Kok, Y. M. Shcherbakova, T. P. C. Schlösser, P. R. Seevinck, T. A. van der Velden, R. M. Castelein, K. Ito, and B. van Rietbergen, “Automatic generation of subject-specific finite element models of the spine from magnetic resonance images,” Frontiers in Bioengineering and Biotechnology, vol. Volume 11 - 2023, 2023.

[35] K. Nispel, T. Lerchl, G. Gruber, H. Moeller, R. Graf, V. Senner, and J. S. Kirschke, “From MRI to FEM: an automated pipeline for biomechanical simulations of vertebrae and intervertebral discs,” Frontiers in Bioengineering and Biotechnology, vol. Volume 12-2024, 2025.

[36] L. Carpenedo, D. Ignasiak, R. Remus, and L. La Barbera, “Advances in musculoskeletal modeling of the thoraco-lumbar spine: A comprehensive systematic review: L. carpenedo et al.,” Annals of Biomedical Engineering, vol. 53, no. 11, pp. 2883–2910, 2025.

[37] G. Holzapfel, T. Gasser, and R. Ogden, “A new constitutive framework for arterial wall mechanics and a comparative study of material models,” Journal of Elasticity, vol. 61, no. 1-3, pp. 1–48, 2000.

[38] D. Nolan, A. Gower, M. Destrade, R. Ogden, and J. McGarry, “A robust anisotropic hyperelastic formulation for the modelling of soft tissue,” Journal of the Mechanical Behavior of Biomedical Materials, vol. 39, pp. 48–60, 2014.

[39] L. D. Landau and E. M. Lifshitz, Theory of Elasticity, vol. 7 of Course of Theoretical Physics. Oxford: Pergamon Press, 3rd ed., 1986.

[40] A. Rohlmann, T. Zander, M. Rao, and G. Bergmann, “Realistic loading conditions for upper body bending,” Journal of Biomechanics, vol. 42, no. 7, pp. 884–890, 2009.

[41] O. C. Zienkiewicz and R. L. Taylor, The finite element method for solid and structural mechanics. Elsevier, 2005.

[42] M. S. Alnæs, J. Blechta, J. Hake, A. Johansson, B. Kehlet, A. Logg, C. Richardson, J. Ring, M. E. Rognes, and G. N. Wells, “The FEniCS Project Version 1.5,” Archive of Numerical Software, vol. 3, 2015.

[43] M. S. Alnaes, A. Logg, K. B. Ølgaard, M. E. Rognes, and G. N. Wells, “Unified form language: A domain-specific language for weak formulations of partial differential equations,” ACM Transactions on Mathematical Software, vol. 40, 2014.

[44] P. Amestoy, I. S. Duff, J. Koster, and J.-Y. L’Excellent, “A fully asynchronous multifrontal solver using distributed dynamic scheduling,” SIAM Journal on Matrix Analysis and Applications, vol. 23, no. 1, pp. 15–41, 2001.

[45] R. Levi, M. Mollura, G. Savini, et al., “CT cadaveric dataset for radiomics features stability assessment in lumbar vertebrae,” Scientific Data, vol. 11, p. 366, 2024.

[46] F. Isensee, P. F. Jaeger, S. A. Kohl, J. Petersen, and K. H. Maier-Hein, “nnu-net: a self-configuring method for deep learning-based biomedical image segmentation,” Nature methods, vol. 18, no. 2, pp. 203–211, 2021.

[47] “3D Slicer image computing platform.” https://www.slicer.org/, 2025. Accessed: February - September 2025.

[48] A. Fedorov, R. Beichel, J. Kalpathy-Cramer, J. Finet, J.-C. Fillion-Robin, S. Pujol, C. Bauer, D. Jennings, F. Fennessy, M. Sonka, J. Buatti, S. Aylward, J. V. Miller, S. Pieper, and R. Kikinis, “3d slicer as an image computing platform for the quantitative imaging network,” Magnetic Resonance Imaging, vol. 30, no. 9, pp. 1323–1341, 2012.

[49] I. Wolf, M. Vetter, I. Wegner, T. Böttger, M. Nolden, M. Schöbinger, M. Hastenteufel, T. Kunert, and H.-P. Meinzer, “The medical imaging interaction toolkit,” Medical image analysis, vol. 9, no. 6, pp. 594–604, 2005.

[50] M. Nolden, S. Zelzer, A. Seitel, D. Wald, M. Müller, A. M. Franz, D. Maleike, M. Fangerau, M. Baumhauer, L. Maier-Hein, et al., “The medical imaging interaction toolkit: challenges and advances: 10 years of open-source development,” International journal of computer assisted radiology and surgery, vol. 8, no. 4, pp. 607–620, 2013.

[51] L. Antiga, M. Piccinelli, L. Botti, B. Ene-Iordache, A. Remuzzi, and D. A. Steinman, “An image-based modeling framework for patient-specific computational hemodynamics,” Medical & biological engineering & computing, vol. 46, no. 11, pp. 1097–1112, 2008.

[52] M. Fedele and A. Quarteroni, “Polygonal surface processing and mesh generation tools for the numerical simulation of the cardiac function,” International Journal for Numerical Methods in Biomedical Engineering, vol. 37, no. 4, p. e3435, 2021.

[53] N. Bogduk, Clinical anatomy of the lumbar spine and sacrum. Elsevier Health Sciences, 2005.

[54] C. Geuzaine and J.-F. Remacle, “Gmsh: A 3-D Finite Element Mesh Generator with Built-in Pre- and Post-Processing Facilities,” International Journal for Numerical Methods in Engineering, vol. 79, pp. 1309–1331, 09 2009.

[55] N. Newell, J. Little, A. Christou, M. Adams, C. Adam, and S. Masouros, “Biomechanics of the human intervertebral disc: A review of testing techniques and results,” Journal of the Mechanical Behavior of Biomedical Materials, vol. 69, pp. 420–434, 2017.

[56] M. Dreischarf, A. Rohlmann, G. Bergmann, and T. Zander, “Optimised in vitro applicable loads for the simulation of lateral bending in the lumbar spine,” Medical engineering & physics, vol. 34, no. 6, pp. 777–780, 2012.

[57] M. Dreischarf, A. Rohlmann, G. Bergmann, and T. Zander, “Optimised loads for the simulation of axial rotation in the lumbar spine,” Journal of Biomechanics, vol. 44, no. 12, pp. 2323–2327, 2011.

